# Dynamic extracellular interactions with AMPA receptors

**DOI:** 10.1101/2025.07.11.664166

**Authors:** Hana Goldschmidt Merrion, Casey N. Barber, Santosh S. Renuse, Jevon Cutler, Simion Kreimer, Alexei M. Bygrave, David J. Meyers, W. Dylan Hale, Akhilesh Pandey, Richard L. Huganir

**Affiliations:** Solomon H. Snyder Department of Neuroscience, The Johns Hopkins University School of Medicine, 725 North Wolfe Street, Baltimore, MD 21205, USA; McKusick-Nathans Institute of Genetic Medicine, Johns Hopkins University School of Medicine, Baltimore, Maryland 21205, United States; Department of Biological Chemistry, The Johns Hopkins University School of Medicine, 725 North Wolfe Street, Baltimore, MD 21205, USA; Department of Pharmacology and Molecular Sciences, The Johns Hopkins University School of Medicine, 725 North Wolfe Street, Baltimore, MD 21205, USA; Department of Laboratory Medicine and Pathology, Mayo Clinic, 200 First Street SW, Rochester, Minnesota 55905, United States

**Keywords:** IgLON, synaptic plasticity, AMPA receptor, proximity proteomics, extracellular

## Abstract

Synaptic plasticity in the central nervous system enables the encoding, storing, and integrating new information. AMPA-type glutamate receptors (AMPARs) are ligand-gated ion channels that mediate most fast excitatory synaptic transmission in the brain, and plasticity of AMPARs signaling underlies the long-lasting changes in synaptic efficacy and strength important for learning and memory.^1,2^ Recent work has indicated that the enigmatic N-terminal domain (NTD) of AMPARs may be a critical regulator of synaptic targeting and plasticity of AMPARs. However, few synaptic proteins have been identified that regulate AMPAR plasticity through interactions with AMPAR NTDs. Moreover, the scope of AMPAR NTD interactors that are important for synaptic plasticity remains unknown. Here, we present the dynamic, extracellular interactome for AMPARs during synaptic plasticity. Using surface-restricted proximity labeling and BioSITe-based proteomics, we identified 70 proteins that were differentially labeled by APEX2-tagged AMPARs after induction of chemical Long-term potentiation of synapses (cLTP) in cultured neurons. Included in this list, were four members of the IgLON family of GPI-anchored proteins (Ntm, OBCAM/Opcml, Negr1, Lsamp). We show OBCAM and NTM directly interact with the extracellular domains of AMPARs. Moreover, overexpression of NTM significantly attenuates the mobility of surface AMPARs in dendritic spines. These data represent a significant first step at uncovering the unexplored extracellular regulation of AMPARs, with broad implications for synapse function and synaptic plasticity.

**Significance Statement:** Over the past 30 years, significant effort has been focused on understanding the mechanisms that induce long-lasting changes in synapse strength (synaptic plasticity) that drive learning and memory. While many studies have investigated intracellular mechanisms that enable plasticity, especially those acting on AMPA-type glutamate receptors (AMPARs), significantly less is known regarding extracellular mechanisms that shape changes in synapse function. Here, we identified 70 proteins that differentially associate with the extracellular region of AMPARs during chemically-induced synaptic plasticity. We show that OBCAM and NTM directly interact with the NTD of AMPARs and regulate their mobility on the surface of neurons. These data advance our understanding of extracellular AMPAR regulation, with broad implications for synapse function and synaptic plasticity.

## Introduction

Information processing in the brain occurs at specialized cellular junctions known as synapses. Transmission at synapses occurs through the pre-synaptic release of neurotransmitters and the post-synaptic sensation of neurotransmitters by neurotransmitter receptors, which convert the chemical neurotransmitter signal into an electrical signal at the post-synaptic membrane. Synapses undergo dynamic reorganization in response to experience in a process known as synaptic plasticity.

Synaptic plasticity can produce long-lasting changes in the efficacy of synaptic transmission, and these changes are a likely physiological correlates of learning and memory. AMPA (alpha-amino-3-hydroxyl-5-methyl-isoxazole-4-propionic acid)-type glutamate receptors (AMPARs) mediate the majority of fast excitatory synaptic transmission throughout the brain and act as effectors of forms of synaptic plasticity^1,2^. The number and organization of AMPARs at synapses change during synaptic plasticity, but the mechanisms by which AMPARs are regulated at synapses at baseline and following plasticity are poorly understood. Understanding the mechanisms by which AMPAR organization is regulated at synapses following plasticity is, therefore, a major focus of synaptic neuroscience.

AMPARs are tetrameric ligand-gated ion channels comprised of four homologous subunits, GluA1-A4^3^. Each subunit possesses four-distinct domains, an amino-terminal domain (NTD) of unknown function, a ligand-binding domain (LBD) specialized for binding the neurotransmitter glutamate, a membrane-embedded transmembrane domain (TMD) which forms the ion channel, and a cytosolic (C)-terminal domain (CTD) which is the site of various post-translational modifications^4–6^ and interacts with intracellular scaffolding proteins^7–11^. Recent work has highlighted a potential role for the understudied NTD in synaptic anchoring and AMPAR plasticity^12–15^. Extending 13 nm into the synaptic cleft, the large, conformationally heterogeneous and sequence-diverse NTD belonging to each AMPAR subunit is optimally positioned to dynamically and selectively interact with proteins in the synaptic cleft^13^. There is growing evidence that interactions with the NTD are functionally significant and play roles in modulating the trafficking, postsynaptic clustering, and ion channel properties of AMPARs^12,14–21^. These lines of evidence suggest that the role of the NTD may be to stabilize AMPAR localization at synapses through direct interaction with other proteins in the synapse. Nevertheless, only a handful of proteins have been shown to engage directly with the AMPAR NTD, including the neuronal pentraxins^19^, Noelins^22^, and recently the NP65 protein^18^. Furthermore, it’s unclear how these protein interactions may change following experience to regulate AMPAR plasticity.

Previous proteomic investigations of AMPAR synaptic interacting proteins have isolated interacting proteins through antibody-mediated pull-down of AMPARs from the brain, followed by mass-spectrometry to identify proteins of interest^23–29^. These approaches have a few caveats. First, AMPARs are transmembrane proteins, and therefore, the site of the putative interaction between an AMPAR and a candidate interactor may be intracellular, extracellular, or intramembrane. Second, AMPARs are often connected to vast intracellular scaffolds within the post-synapse, so interactors identified with affinity chromatography may lack a specific direct interaction with AMPARs^8–11,30^. This is especially relevant given that AMPARs are exquisitely organized within the post-synapse at spatially restricted sites^31–37^. Finally, traditional affinity chromatography techniques favor the detection of stable, high-affinity interactions and may miss important transient or low-affinity interactions that are disrupted by biochemical isolation.

We sought to test the hypothesis that the extracellular domains of AMPARs participate in dynamic protein-protein interactions that facilitate AMPAR-mediated synaptic plasticity using a surface-restricted proximity-labeling approach^38–40^. We fused genetically encoded ascorbate peroxidase (APEX2) to the NTD of recombinant AMPARs and used proximity biotinylation to systematically identify protein interactions with the extracellular region of AMPARs in cultured neurons before and after chemical induction of synaptic plasticity (cLTP). We successfully identified 70 proteins that were differentially labeled by APEX2-tagged AMPARs after cLTP. Included in this list were four members of the IgLON family of GPI-anchored proteins (Ntm, OBCAM/Opcml, Negr1/Kilon, Lsamp)^41–43^. Biochemical analysis shows OBCAM and NTM directly interact with the extracellular region of AMPARs. Functional studies further implicate NTM in the regulation of surface AMPAR mobility in spines. While the precise roles of IgLONs and other proteins identified in our screen in AMPAR-mediated plasticity have yet to be investigated, these data show that the extracellular region of AMPARs does indeed dynamically engage with a myriad of proteins on the surface of neurons and impacts AMPAR mobility.

## Results

### A Proximity Biotinylation Strategy for Capturing Dynamic, Extracellular AMPAR Interactions

To identify the full complement of extracellular proteins that regulate AMPARs during synaptic plasticity, we designed an unbiased, proteomics-based approach using APEX2 proximity labeling. APEX2 is a small, monomeric peroxidase reporter that catalyzes the oxidation of biotin-phenol (BP) in the presence of H_2_O_2_ (**Fig. 1A**). The resulting biotin-phenoxyl radicals are short-lived (<1 msec), membrane impermeable, and highly reactive species that covalently attach to electron-rich amino acids on proteins located <20 nm from the APEX2 active site^38,39^. This allows for acute, permanent labeling of proteins immediately proximal to AMPARs. To enhance the selectivity of our approach to identify extracellular AMPAR interacting partners, we synthesized a membrane impermeant variant of BP (BxxP)^38^. Biotinylated proteins labeled with BxxP during a one-minute reaction are then readily purified from cells and identified by mass spectrometry (MS). The temporally controlled, spatially restricted, and covalent labeling by APEX2 in live cells makes this an ideal system for identifying transient and dynamic interactions with extracellular domains of AMPARs.

**Figure 1.**
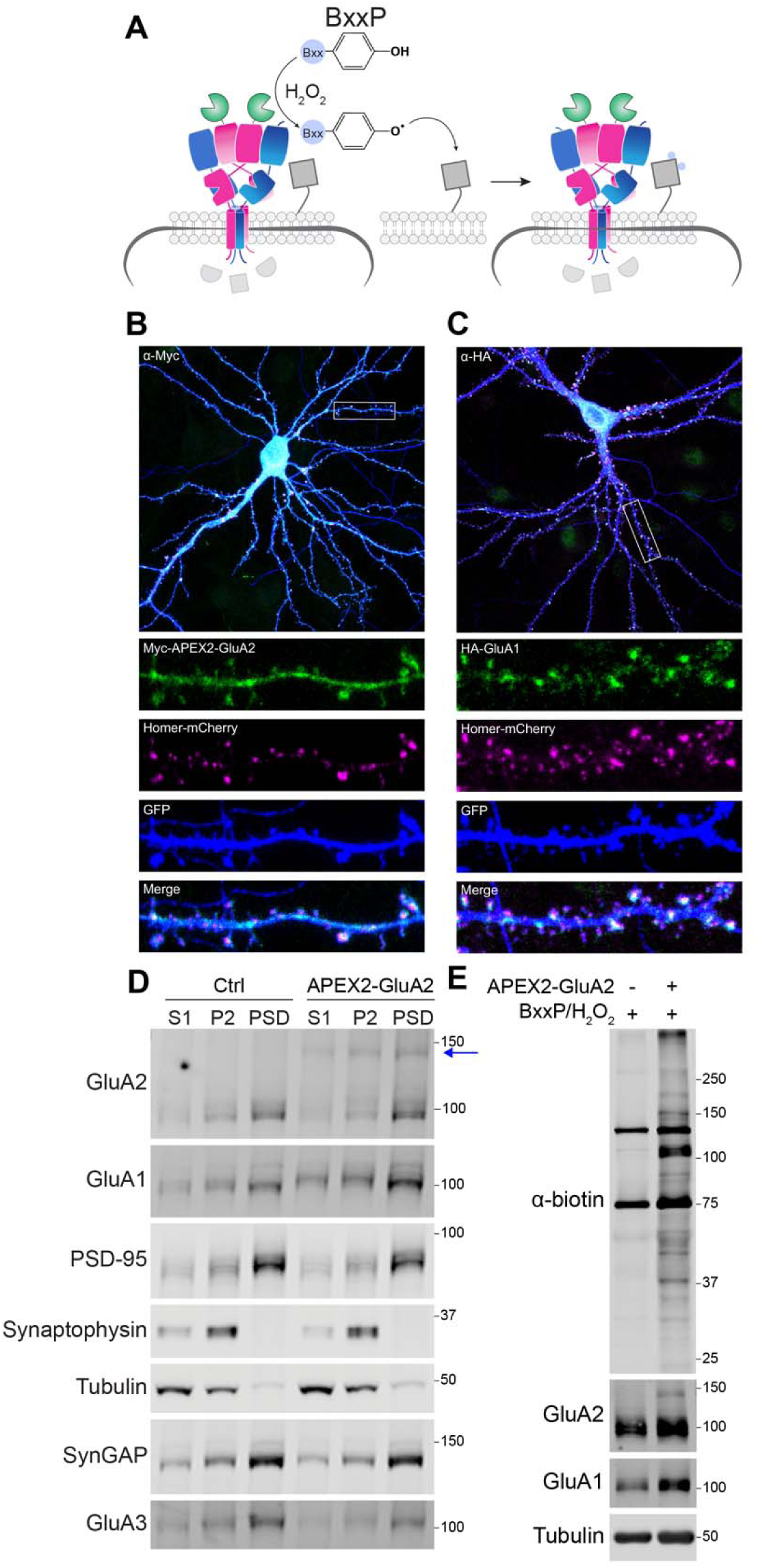
Surface-restricted proximity labeling with APEX2-tagged AMPA receptors in neurons. **(A)** Schematic of APEX labeling in neurons using APEX2-tagged AMPARs. Labeling by APEX2-AMPARs (APEX2-GluA2, pink, HA-GluA1, blue) is initiated by adding membrane impermeant BxxP to live neurons for 1 minute together with H2O2. APEX2 converts BxxP into a phenoxyl radical which covalently attaches to tyrosine residues on the extracellular regions of nearby proteins (eg. blue circles on grey square protein). Intracellular interacting proteins are not labeled (light gray) as phenoxyl radicals cannot cross the membrane. **(B-C)** APEX2-tagged AMPARs localize to excitatory synapses. Cultured hippocampal neurons transfected with APEX2-GluA2, HA-GluA1 and homer mCherry were immunostained with antibodies against Myc (B, APEX2-GluA2) or HA (C, HA-GluA1) along with mCherry and GFP. Both APEX2-AMPAR subunits co-localize with homer-mCherry in dendritic spines. **(D)** Western blot analysis of subcellular fractions purified from cortical neurons expressing APEX2-AMPARs. The band corresponding to APEX2-GluA2 (150kDa band) is enriched in crude membrane (P2) and postsynaptic density (PSD) fractions compared to the crude cytosol (S1), similar to endogenous GluA2 (100kDa). Fractionation efficiency and purity is demonstrated by immunoblots of synaptic proteins (PSD-95, synaptophysin, GluA3, SynGAP) and a cytosolic protein (tubulin). **(E)** Western blot analysis of APEX labeling in neurons. Cortical neurons expressing APEX2-AMPARs were labeled for 1 minute with 300uM BxxP/1mM H_2_O2 and whole-cell lysates were harvested after the reaction was quenched. Immunoblotting with anti-biotin antibodies shows robust biotinylation only when APEX2-AMPARs are expressed. Labeling reactions performed in control neurons (no APEX2 expression) did not show any additional biotinylation above endogenous biotinylated proteins (75kDa, 140kDa).

To enable the selective tagging of extracellular AMPAR interactions we generated a N-terminal, extracellular-facing fusion of APEX2 with the GluA2 subunit (Myc-APEX2-GluA2) which was co-expressed in primary rat neurons with HA-tagged GluA1 (HA-GluA1, **Fig. 1A**). This cloning and expression strategy has been previously used to successfully express N-terminally-tagged, GluA1/2 AMPARs that traffic and function similar to wildtype receptors in vitro and in vivo^44–47^.

We first verified that APEX2-GluA2/HA-GluA1 AMPARs (APEX2-AMPARs) are properly trafficked to synapses in intact neurons. For these experiments, APEX2-AMPARs were co-expressed via transfection in rat hippocampal neurons with the postsynaptic protein, mCherry-tagged Homer, along with a cell fill (GFP) to visualize neuronal morphology. Immunolabeling of APEX2-AMPARs with antibodies against either the Myc-(APEX-GluA2) or HA-(GluA1) tagged subunits showed a punctate distribution along dendrites and within dendritic spines (**Fig. 1B-C**). Moreover, both APEX-AMPAR subunits colocalized with the Homer-mCherry in dendritic spines, demonstrating that APEX2-AMPARs correctly traffic to excitatory synapses in mature neurons.

We further verified these data using biochemical fractionation of mature rat cortical neurons expressing APEX2-AMPARs. Western blot analysis using an antibody specific for GluA2 showed both endogenous and APEX2-tagged subunits (∼100 kDa and ∼140 kDa bands, respectively) are similarly enriched in membrane (P2) and postsynaptic density (PSD) fractions from control and APEX2-expressing neurons (**Fig. 1D**). APEX2-AMPARs expression was titrated to ensure that both subunits were expressed stably and synaptically localized for at least 3 weeks in culture. Additional immunoblotting for neuronal and synaptic proteins, including PSD-95, synaptophysin, tubulin, SynGAP, and GluA3, showed no significant differences in expression or subcellular distribution between control and APEX2-AMPAR-expressing neurons (**Fig. 1D**). Taken together, these data show that APEX2-AMPARs are stably expressed in neurons and properly trafficked to excitatory synapses.

To selectively identify surface-expressed AMPAR interactors, we synthesized the membrane impermeant biotin-phenol substrate (BxxP) to use for APEX2-dependent biotinylation previously used to map surface proteomes^38^. To evaluate the labeling activity with BxxP, neurons expressing APEX2-AMPARs or control neurons were labeled live for 1 minute with BxxP in the presence of H_2_O_2_ and subsequently lysed. Western blot analysis with an anti-biotin antibody shows robust biotinylation when all three components required for APEX-dependent biotinylation are present: APEX2, H_2_O_2_ and BxxP (**Fig. 1E).** Only endogenous biotinylated proteins (bands at 140 and 75 kDa) are observed when control neurons, not expressing APEX2-AMPARs, were labeled (**Fig. 1E**). Taken together, APEX2-AMPARs are properly trafficked to synapses, are stably expressed for three weeks in neurons and demonstrate specific and temporally controlled BxxP labeling of endogenous proteins.

### Identification of Dynamic Extracellular AMPAR Interactions during cLTP

We predicted that following the induction of cLTP, interactions with the extracellular region of synaptic AMPAR would be modified to facilitate changes in synapse strength. If this is the case, the BxxP-biotinylation of extracellular interactors would change between control and stimulated conditions due to their proximity with the AMPAR, where the gain or loss of protein interactions would be reflected by an increase or decrease in biotinylation, respectively (**Fig. 2A**).

**Figure 2.**
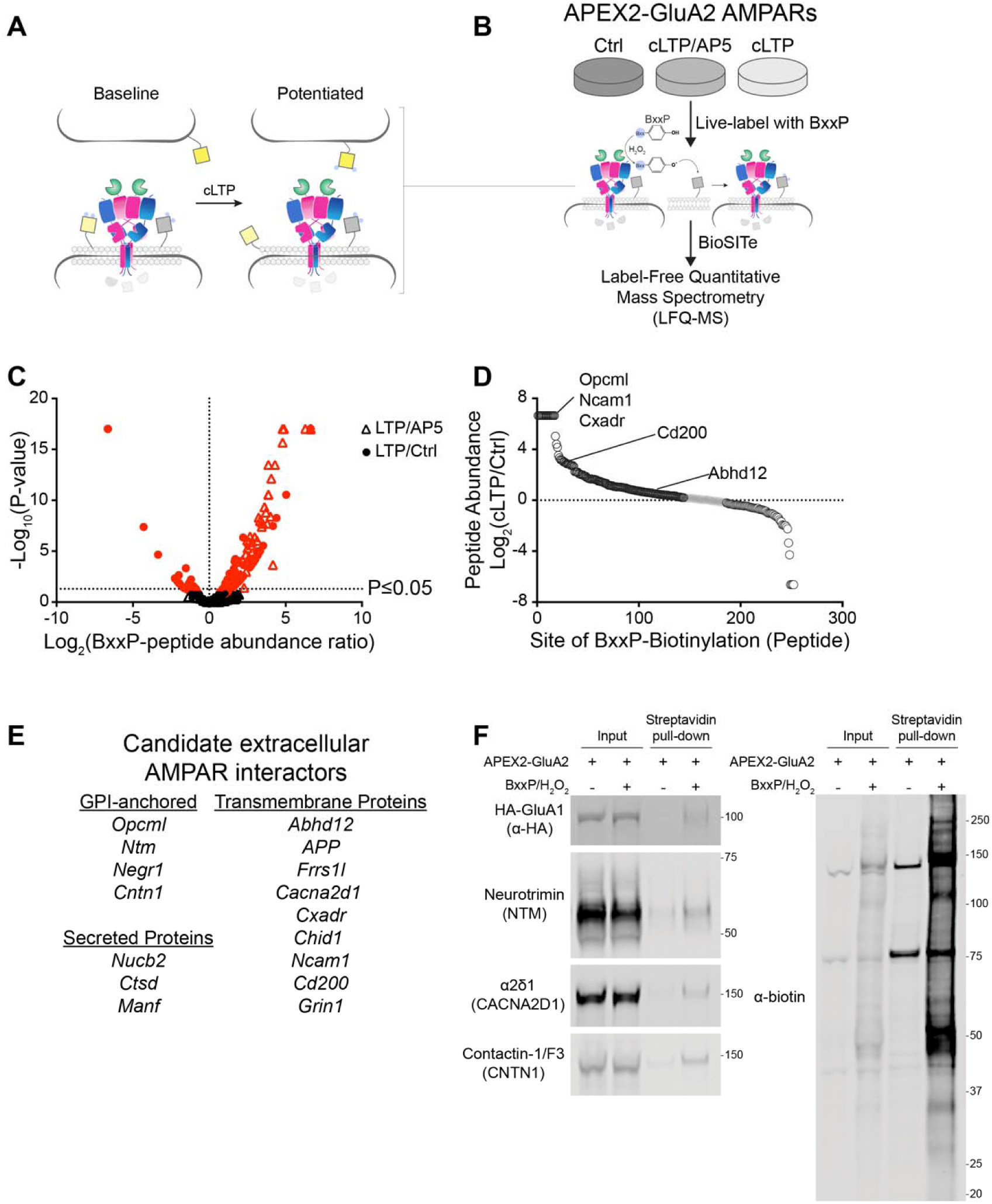
Differential biotinylation by APEX2-GluA2 AMPARs after cLTP. **(A)** Schematic of predicted differential labeling by APEX2-AMPARs before and after cLTP induction. **(B)** Schematic of experimental design of BioSITe-based identification of dynamic interactions with the extracellular region of AMPARs during cLTP. Three independent BioSITe-based proteomic experiments were performed. **(C)** Volcano plot of differential biotinylation by APEX2-AMPARs after cLTP. Peptides that significantly changed in abundance after cLTP are shown in red (P≤0.05). **(D)** Quantification of the change in biotinylation following cLTP compared to control (Y-axis) is shown for individual BxxP-peptides identified by BioSITe-proteomics (X-axis). Examples of individual peptides only identified in cLTP samples but not the control are shown for Opcml, Ncam1 and Cxadr. **(E)** Candidate, extracellular AMPAR interactors identified by BioSITe-MS that were selected for subsequent analysis. **(F)** Western blot analysis of streptavidin-enriched whole-cell lysates after APEX labeling by APEX2-AMPARs in neurons. Immunoblotting with antibodies against candidate proteins NTM, CACNA2d1 and CNTN1 detects bands in streptavidin-enriched fractions only when cells were labeled, confirming BioSITe-MS data. HA-GluA1 immunoblot is shown as a positive control for APEX labeling.

To capture dynamic extracellular interactions during synaptic plasticity in live cells, we set up three conditions for APEX labeling in neurons (**Fig. 2B**). Neurons were either (1) mock stimulated (Ctrl) or (2) stimulated with a chemical cocktail that induces an LTP-like phenomenon by activating endogenous NMDARs with glycine (cLTP^45^). For our third condition, we included an additional control for the stimulation where cLTP was induced in neurons in the presence of the NMDA receptor inhibitor, AP5 (cLTP/AP5). Mature rat cortical neurons (DIV 19) were treated with each of these conditions, and live cells were subsequently labeled for one minute with BxxP and harvested for quantification and analysis using BioSITe-based proteomics.^17^ The BioSITe-based approach enables the direct detection of BxxP-biotinylated peptides by mass spectrometry (MS), a feature greatly limited by conventional streptavidin-based methods^48–51^.

To accurately identify proteins that were differentially biotinylated by APEX2-AMPARs following cLTP induction, we performed three independent proteomic experiments, each containing biological replicates for control and stimulated samples (**Fig. 2A-B**). Using this approach, we identified >500 BxxP-biotinylated peptides that could be quantified in cLTP/Ctrl or cLTP/AP5 conditions (**Supplemental Table 1, Supplemental Table 2**). These peptides mapped to >140 different proteins, with 97.9% overlap of proteins quantified in the two conditions (**Supplemental Fig. 1A**).

We further analyzed differential BxxP-biotinylation across biological and experimental replicates as ratios across the three conditions (cLTP/Ctrl, cLTP/AP5, Ctrl/AP5) to identify specific interactions as well as those that significantly increase after cLTP induction. To do this, we applied two filters to the data to remove non-specific interactors and false-positive changes in biotinylation (BxxP-peptide increase ≥1.2-fold in Ctrl/AP5, **Supplemental Fig. 1B-C, Supplemental Table 3, Supplemental Table 4**).

Our final list of candidate extracellular AMPAR interactors, we identified proteins containing BxxP-peptides that were differentially biotinylated after cLTP induction compared to controls (**Fig. 2C-D**, **Supplemental Table 3**). As expected, included in this list were peptides mapping to the extracellular domains of AMPAR subunits (GluA1-3) and AMPAR auxiliary subunits (FRRS1l, ABHD12).

We were particularly interested in proteins containing BxxP-peptides that significantly increased after cLTP induction as they might reflect the selective, activity-dependent association with the extracellular region of GluA1/2 AMPARs. Candidate interactors prioritized for further characterization and analyses were proteins containing ≥1 BxxP-peptide that significantly increased after cLTP compared to controls, as well as proteins implicated in AMPAR and/or synapse function. Using these criteria, we created an initial list of candidate AMPAR interactors, which included secreted proteins, GPI-anchored proteins, and receptors/transmembrane proteins (**Fig. 2E**).

We first confirmed that identified candidate proteins by BioSITe-MS are biotinylated by APEX2-AMPARs in neurons using an alternative approach. For these experiments, neurons expressing APEX2-AMPARs were labeled with BxxP and biotinylated proteins were isolated from whole-cell lysates using streptavidin beads. Western blotting analysis using antibodies specific to candidate proteins NTM, CACNA2D1, and CNTN1 shows they are selectively enriched by streptavidin only in lysates where APEX labeling occurred (**Fig. 2F)**. Blotting for HA-GluA1 served as a positive control for BxxP-biotinylation by APEX2-AMPARs.

### IgLONs interact with the extracellular region of AMPA receptors

One family of proteins that stood out in the BioSITe-proteomic data were the <u>Ig</u>-superfamily <u>L</u>AMP, <u>O</u>BCAM/OPCML, and <u>N</u>eurotrimin (IgLONs). This family of Ig-like, GPI-anchored cell adhesion molecules consists of five proteins: OBCAM, NTM, Negr1/Kilon, LSAMP, IgLON5. IgLON proteins are known to homo- and heterodimerize with high-affinity in neurons (**Supplemental Fig. 2A**) and have been implicated in neurite outgrowth, axon fasciculation and synapse formation as well as several neurological and psychiatric disorders^41–43,52–55^. Recent data suggests that individual IgLONs harbor distinct functions in mature neurons to those identified in the developing nervous system and might play key roles in the establishment and maintenance of specific neural circuits^53,55^. Importantly, four of the five IgLONs were found to be differentially biotinylated by APEX2-AMPARs after cLTP using BioSITe-based proteomics (**Fig. 3A, Supplemental Table 1**).

**Figure 3.**
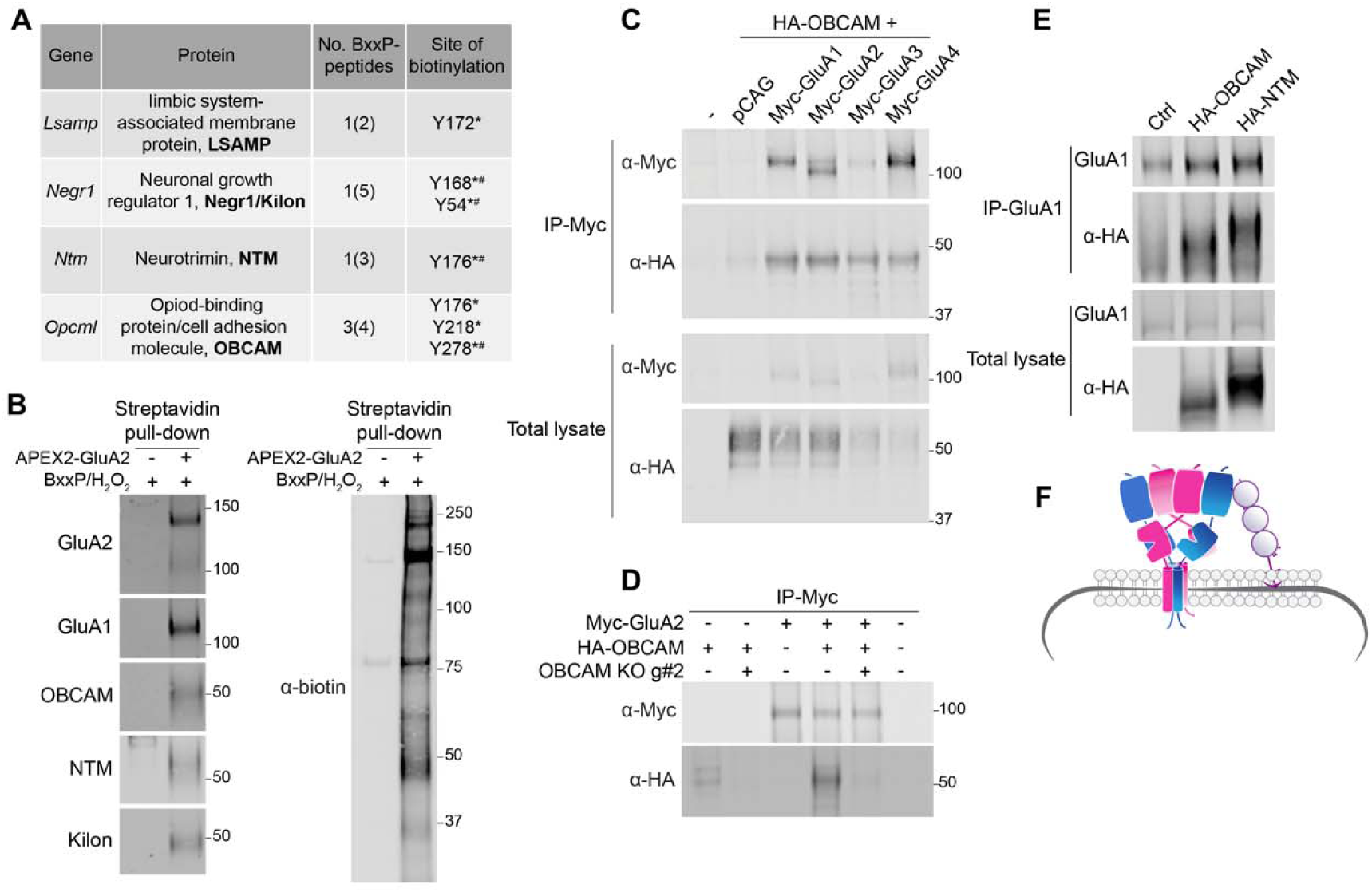
IgLONs interact with the extracellular region of AMPARs. **(A)** Differential biotinylation of IgLON proteins by APEX2-AMPARs identified by BioSITe-MS. Four Iglons identified are listed. Number of unique peptides identified and total number of BxxP-modified peptides (No. shown in parentheses) are shown for each gene, as well as the site of biotinylation (tyrosine, Y). *LTP/Ctrl, P≤0.05, # LTP/AP5 P≤0.05. **(B)** Western blot analysis of streptavidin-enriched lysates after APEX labeling by APEX2-AMPARs in neurons. **Left,** Immunoblotting with antibodies against IgLON proteins OBCAM, NTM and Kilon/NEGR1 detects bands in streptavidin-enriched fractions only in samples where APEX2-AMPARs were expressed (lane 2). No IgLONs were detected in fractions isolated from control labeled neurons where APEX2 was not expressed (lane 1). Immunoblots for GluA2 and GluA1 are shown as positive controls. **Right,** immunoblotting with anti-biotin antibodies shows robust APEX labeling only when APEX2-AMPARs are expressed (right two lanes). **(C)** AMPARs interact with OBCAM in HEK cells. HA-tagged OBCAM was expressed in HEK cells along with individual, Myc-tagged AMPAR subunits (GluA1-4). AMPAR complexes were immunoprecipitated (co-IP) from whole-cell lysates using anti-myc antibodies. Immunoblotting with anti-HA antibodies show OBCAM is present in AMPAR precipitates, but not in controls (wild-type or cells empty pCAG vector). Immunoblotting of whole-cell lysates (lower panels) shows relative expression of each AMPAR subunit and OBCAM prior to co-IP. HA-NTM and HA-Kilon were also co-immunoprecipitated by Myc-AMPARs (data not shown). **(D)** OBCAM directly interacts with GluA2-containing AMPARs. Co-IP experiments in HEK cells show HA-OBCAM is present in GluA2 precipitates (lane 4). To demonstrate specificity of this interaction we used a CRISPR KO guide targeting OBCAM cDNA. Expression of OBCAM is successfully depleted by expression of CRISPR knockout guides targeting the cDNA (OBCAM KO g#2) (lane 2) and thereby depletes OBCAM from GluA2 precipitates (lane 5). **(E)** IgLONs interact with endogenous AMPAR complexes in neurons. Neurons expressing HA-tagged OBCAM or NTM were fractionated and endogenous AMPARs were purified from crude membrane fractions using anti-GluA1 antibodies. Immunoblotting with anti-HA antibodies shows HA-tagged IgLONs precipitate with GluA1 AMPARs, but not in control neurons. HA-IgLONs were found in AMPAR complexes isolated using anti-GluA2 antibodies specific for the N- and C-terminus (Supplemental Figure 2C-D)**. (F)** Schematic of IgLON-AMPAR complexes. IgLONs are GPI-anchored proteins with all 3 protein domains expressed on the cell surface. Picture shows IgLONs interacting with AMPARs in cis (on the same side of the membrane). It is possible that they interact with AMPARs in cis or trans (across the synapse/from another membrane).

Together, these data implicate the association of IgLONs with AMPAR complexes in neurons. To test this, we first confirmed that IgLON proteins identified by BioSITe-MS are biotinylated by APEX2-AMPARs in neurons using streptavidin capture and western blotting. As shown in **Figure 3B**, OBCAM, NTM and Kilon/NEGR1 are selectively enriched by streptavidin beads when BxxP-biotinylation by APEX2-AMPARs occurred.

To determine if IgLONs associate with AMPAR complexes, we used heterologous cells to evaluate the association between HA-tagged IgLONs and Myc-tagged AMPARs using co-immunoprecipitation. OBCAM co-immunoprecipitated (co-IP) with all four AMPAR subunits and was not found in controls (**Fig. 3C**). We further interrogated the interaction with GluA2, showing OBCAM and NTM were found in GluA2 precipitates, demonstrating a direct interaction between IgLONs and AMPARs in HEK cells (**Fig. 3D, Supplemental Fig. 2B**). When CRISPR knockout (KO) guides targeting OBCAM or NTM were co-expressed in this system, both IgLONs were absent from GluA2 precipitates. These data demonstrate that OBCAM and NTM directly interact with GluA2-containing AMPARs.

We next tested if IgLONs interact with AMPAR complexes in neurons. For these experiments, HA-tagged IgLONs were expressed in neurons and endogenous AMPAR complexes were purified from neuronal membrane fractions. HA-OBCAM and HA-NTM co-immunoprecipitated with endogenous GluA1-containing AMPARs isolated using antibodies against the GluA1 C-terminus (**Fig. 3E**). HA-IgLONs also co-immunoprecipitated with GluA2-containing AMPARs that were isolated using antibodies against the N- or C-terminus of GluA2 (**Supplemental Fig. 2E-F**). Together, these data demonstrate that IgLONs associate with AMPARs in neurons (**Fig. 3F**).

Synaptic AMPAR trafficking is critical to synapse function and plasticity. If IgLONs are involved in regulating synaptic AMPARs we predicted that they would be targeted to excitatory synapses. Using biochemical fractionation, we analyzed the subcellar distribution of endogenous IgLONs in neurons. Western blotting of synaptic fractions isolated from mature cortical neurons found both endogenous OBCAM and NTM proteins are significantly enriched in membrane and PSD fractions (**Fig. 4A-B**). These data are consistent with reports of the synaptic localization of OBCAM and NTM^42,54^. Next, we used immunocytochemistry to further examine the subcellular distribution of tagged IgLON proteins, OBCAM, and NTM transfected into cultured rat hippocampal neurons. Both HA-tagged IgLONs showed a punctate distribution throughout the soma as well as along dendrites and axons (**Supplemental Fig. 3**). Importantly, both IgLONs co-localized with Homer-mCherry along dendrites and within dendritic spines, showing that recombinant IgLONs localize to excitatory synapses (**Fig. 4C-D).** Collectively, these data show OBCAM and NTM are localized to the PSD of excitatory synapses and, therefore, implicate these IgLONs in regulating synaptic AMPAR function.

**Figure 4.**
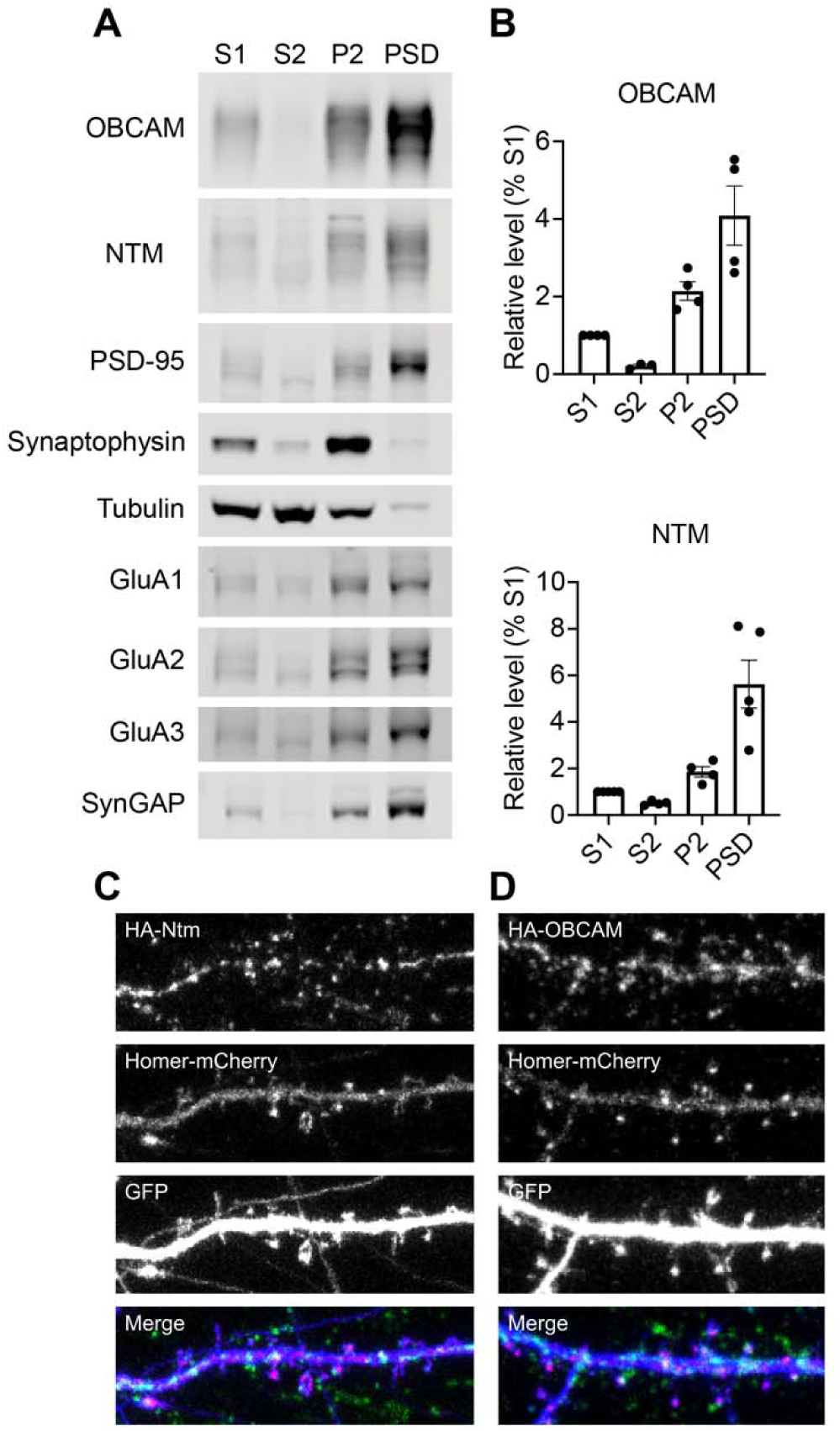
IgLONs localize to excitatory post-synapses. **(A)** Western blot analysis of subcellular fractions purified from cortical neurons. Immunoblotting with antibodies again OBCAM and NTM show both IgLONs are enriched in crude membrane (P2) and postsynaptic density (PSD) fractions compared to the crude lysate (S1) and cytosolic (S2) fractions. Fractionation efficiency and purity is demonstrated by immunoblots of synaptic proteins (PSD-95, synaptophysin, GluA1-3, SynGAP) and a cytosolic protein (tubulin). *N=4-5 experimental replicates, N=3-5 replicates for each fraction.* 6µg of each fraction loaded. **(B)** Quantification of relative levels of endogenous NTM (top) and OBCAM (bottom) protein in each subcellular fraction. Relative values were normalized to S1 fraction for each experiment. N=3-5**. (C-D)** Distribution of HA-tagged IgLONs along dendrites of primary hippocampal neurons. Cells were transiently transfected with 0.3 µg of HA-IgLON cDNA. Immunostaining with antibodies against HA along with mCherry and GFP shows some IgLON puncta co-localize with homer-mCherry in dendritic spines. In merge, blue = GFP, red = mCherry, green = HA-IgLON.

### IgLONs regulate surface AMPAR mobility

Since IgLON proteins are GPI-anchored into the outer leaflet of the plasma membrane, we hypothesized that they might influence the mobility of surface-expressed AMPARs in neurons. To test this, we enhanced or depleted OBCAM and NTM protein levels in neurons using overexpression plasmids or CRISPR-mediated KO guides (**Supplemental Fig. 4**), respectively. To monitor the dynamics of surface-expressed AMPARs, we added a super ecliptic pHluorin (SEP) tag to the extracellular N-terminus of GluA2. SEP is a pH-sensitive variant of GFP that selectively fluoresces at neutral pH and is quenched at acidic pH^56^. In live cells, SEP-GluA2 reports on the dynamics of functional receptors at the cell surface, as the fluorescence of receptors localized in acidic, intracellular compartments, such as early and recycling endosomes, are quenched^57,58^. To measure AMPAR surface mobility, we used fluorescence recovery after photobleaching (FRAP) to quantify SEP-GluA2 dynamics in spines in live neurons.

To test if IgLONs regulate surface/synaptic AMPAR dynamics, neurons were transiently transfected with SEP-GluA2, Myc-GluA1along with dsRED (to observe neuronal morphology) and either CRISPR KO guides targeted to endogenous IgLONs or overexpression plasmids. Individual spines on transfected neurons were targeted for photobleaching, and unbleached, neighboring spines on the same cell were used as controls for FRAP analysis (**Fig. 5A**). Surface AMPAR mobility at synapses was then assessed by FRAP, which reflects the rate of exchange of bleached SEP-GluA2 AMPARs within individual spines with nonsynaptic unbleached SEP-GluA2 AMPARs.

**Figure 5.**
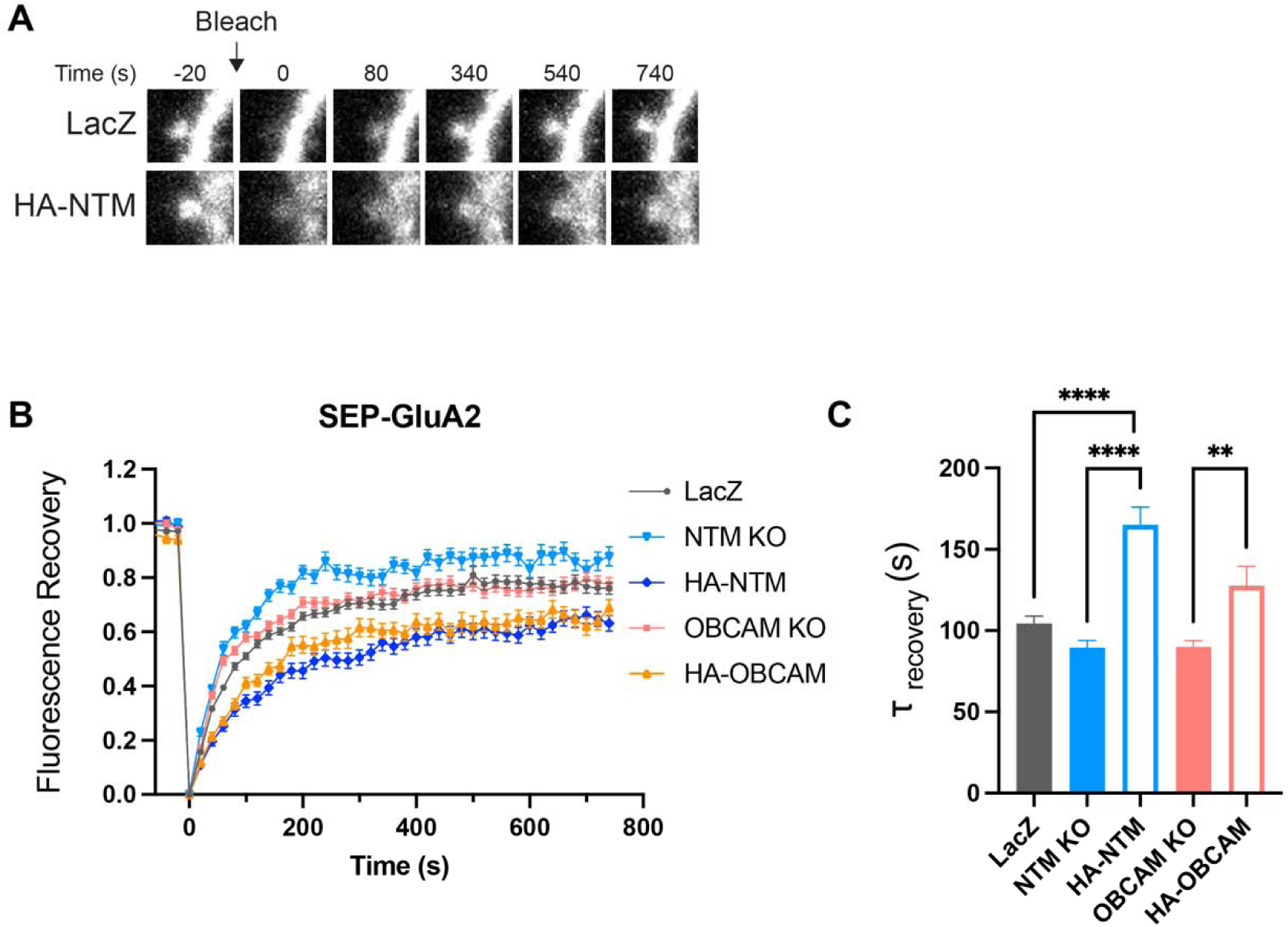
IgLONs stabilize AMPARs at excitatory post-synapses. **(A)** Representative time series of SEP-GluA2 recovery after photobleaching (FRAP). *Top series*: recovery in the control condition when cells were co-transfected with lacZ. *Bottom series:* recovery in the presence of co-transfected NTM. (**B**) Time course of SEP-GluA2 FRAP in cultured rat neurons transfected with control lacZ plasmid (black circles), NTM knockout guide RNA (light blue triangles), NTM overexpression plasmid (dark blue diamonds) OBCAM knockout guide RNA (melon squares), or OBCAM overexpression plasmid (orange triangles). **(C)** Recovery constants for conditions shown in **(B)**. Both NTM overexpression (light blue) and OBCAM overexpression (light orange) significantly slowed SEP-GluA2 FRAP. *One-way ANOVA with multiple comparisons =* α*<0.05*. * = p<0.05; ** = p<0.01; *** = p<0.001; **** = p<0.0001.

Of all the conditions examined, only overexpression of NTM was found to significantly alter the rate of FRAP recovery compared to control neurons (**Fig. 5B-C**, LacZ = 104.4±4.6 sec HA-NTM=165.0±11sec). Similar to NTM, overexpression of OBCAM also attenuated AMPAR mobility, however, it was not statistically significant from control neurons. Loss of either IgLON had the opposite effect on AMPAR mobility, speeding up the recovery rate after photobleaching. However, these effects were small, and while significantly different from overexpression neurons, they were not statistically different from control neurons. Together, these data show that overexpression of NTM significantly slows the mobility of GluA2-containing AMPARs.

## Discussion

Dynamic regulation of AMPARs underlies forms of synaptic plasticity that important for learning and memory. Using unbiased and quantitative global-proteomic analysis of BxxP-biotinylation by APEX2-tagged AMPARs, we identified extracellular interactions with AMPARs that change following plasticity. This approach is advantageous for four principal reasons. First, the 20nm labeling radius of APEX2 ensures that only the proteins most proximal to AMPARs are labeled in our preparation. AMPARs have previously been reported to be organized in nanodomains roughly 70nm in diameter^33,36,37,59^. APEX2 labeling should, therefore, label proteins concentrated within this nanodomain. Second, the reactive species produced by our BxxP labeling is membrane impermeable, which restricts our proteomic analysis to extracellular synaptic cleft constituents. This is reflected by the absence of intracellular scaffolding proteins, like PSD95 (DLG4), in our dataset. Third, the permanent, covalent attachment of the biotin moiety to nearby proteins ensures that low-affinity or transient interactions are not lost during biochemical extraction. Finally, BioSITE proteomics ensures the detection only of proteins that are within the range of APEX2 and not proteins that are bound to AMPARs through an intermediate protein like an intracellular scaffold. For these reasons, our dataset is a unique snapshot of the extracellular AMPAR interactome immediately following a plasticity-inducing stimulus.

Among our candidate plasticity interactors are several proteins that have been identified previously to interact with AMPARs including *Frrs1l*^23,24,27,60^, *Abhd12*^23,24,27^, and most recently *Cacna2d1*^29^. Both *Frrs1l* and *Abhd12* have been proposed to complex with AMPARs in the ER during AMPAR biosynthesis. Deletion of *Cacna2d1* and *Frrs1l* has been reported to impair AMPAR plasticity^29,60^. However, the presence of *Frrs1l* in our dataset suggests that the interaction between *Frrs1l* may persist at the postsynaptic membrane and that the role of *Frrs1l* in regulating AMPAR plasticity may be acute and at synaptic sites. Alternatively, the presence of *Frrs1l* in our dataset may reflect rapid delivery of AMPARs to the postsynaptic membrane following cLTP. In this interpretation, our 15 minute labeling scheme following cLTP captures ER-resident proteins chaperoning AMPARs to the membrane that have not yet had time to fully dissociate from GluA subunits.

Several notable AMPAR interactors are also missing from our dataset, including the Neuronal Pentraxins (NPTXs), which have been previously proposed to regulate AMPAR clustering at synaptic sites^16,17,19,61–63^. The absence of the NPTXs in our dataset and other proteomic datasets^23–28^ targeting AMPARs may indicate that the influence of the NPTXs on synaptic AMPAR dynamics is restricted to sparsely populated neuronal cell types or during more specialized forms of plasticity. For instance, in parvalbumin-expressing interneurons (PV-INs), NPTX2 and NPTXR have been demonstrated to regulate AMPAR content at excitatory synapses on PV-INs during homeostatic scaling^16,17^. The absence of NPTX2 from our dataset may also reflect the temporal specificity of NPTX2 expression following plasticity-inducing stimuli, as previous reports show NPTX2 expression in the dentate gyrus peaking six hours following stimulation^64^. Likewise, LRRTM4, which has been previously identified in AMPAR proteomic screens^23,24,27^ is missing from our data. This may indicate that LRRTM4 is not part of the core AMPAR nanocolumnar assembly and may instead be a product of AMPAR interactions with intracellular scaffolding molecules. Finally, AMPAR auxiliary subunits like TARPs and cornichons, which appear often in AMPAR proteomic datasets^23,24,27,28^, are also absent from our dataset. TARPs possess very limited extracellular domains, only extending a few nanometers from the cell membrane into the synaptic cleft, while cornichons lack extracellular domains entirely^65–68^. The absence of these common AMPAR interacting proteins from our dataset likely reflects the specificity of our labeling scheme for extracellular proteins.

Unique to our dataset is the identification of four IgLON proteins (NTM, OBCAM, Kilon/NEGR1, LSAMP) that were differentially biotinylated by APEX-AMPARs after cLTP. We confirmed that these proteins are indeed biotinylated by APEX-AMPARs in cultured neurons (**Fig. 3**), and further confirmed that APEX-Biotinylation does indeed reflect a direct association between IgLON proteins and AMPARs subunits (**Fig. 3C-E**). IgLON proteins have been previously reported to form homo- and hetero-dimers in the central nervous system^42,43,53^, and have been principally studied in the context of neurite outgrowth^41,43,52^. Structural and biochemical analysis of IgLON homo- and hetero-dimers has suggested that IgLON proteins could be trans-synaptically engaged through their first Immunoglobulin domains^53^. This would make IgLON proteins potential candidates for aligning AMPARs with pre-synaptic release machinery within trans-synaptic nano-columns. However, the relative enrichment of the NTM OBCAM in the PSD relative to P2 in our data (**Fig. 4A,B)** may instead suggest that IgLONs dimerize in *cis* on the post-synaptic membrane rather than bridge the pre- and post-synaptic membranes. Moreover, their punctate distribution showed incomplete overlap with mCherry-Homer (Fig. 4C-D), indicating IgLONs might localize to more than just excitatory synapses.

Until now, the interactions of IgLON proteins with other synaptic components has remained largely unexplored. Based on their established role in cell adhesion and enrichment in the PSD (**Fig. 4B-C**) we speculated that IgLONs might regulate the mobility of surface-expressed AMPARs in spines. Using a FRAP-based imaging approach, we found a modest but significant effect of NTM overexpression on surface AMPAR mobility in dendritic spines (**Fig. 5**). The more modest impact on AMPAR diffusion by NTM or OBCAM deletion likely reflects compensation by the remaining three highly-homologous IgLON family members that were also detected in our screen to be AMPAR interactors. These data indicate that the interaction between IgLONs and AMPARs functionally impacts the retention of AMPARs at synaptic sites and suggests that this may extend to their sub-synaptic organization. Further analysis will be necessary to explore whether IgLON proteins are integrated within AMPAR-containing nanodomains, or whether IgLON proteins exist as a kind of diffusion barrier at the periphery of AMPAR nanodomains within the synapse.

One open question from our study is how the AMPAR interactome changes over time after cLTP stimulation. Super-resolution analysis of synaptic nanodomains following cLTP has revealed that different synaptic proteins change their organization on different time scales following cLTP^37,59^. Both GluA2 and the post-synaptic scaffolding protein PSD95 increased their synaptic content following cLTP, but it has been reported that while PSD95 reached its maximal increase by 30 minutes post-stimulation, GluA2 content at individual spines was still increasing at 120 minutes^59^. It seems likely that different biochemical mechanisms exist at the synapse to coordinate individual aspects of the cLTP response. Our labeling paradigm captured proteins in close proximity to AMPARs in a single snapshot in time 15 minutes after induction of cLTP. In this light, it is interesting to consider whether the candidate interactors in our dataset that increase in biotinylation following cLTP represent stable AMPAR interactors that continuously accumulate following cLTP or whether they are part of a transitional mechanism that signals the switch from the baseline to the potentiated state at the synapse. Future studies should include later time points following cLTP induction to address this question.

In summary, we identified candidate extracellular AMPAR interacting partners that change following a form of synaptic plasticity. Using unbiased, proximity-based proteomics, we further show these interactions are dynamic, which resulted in differential biotinylation patterns before and after synaptic plasticity. We confirmed our mass spectrometry data using biochemical and cell biological approaches, showing a direct interaction between IgLON proteins OBCAM and NTM and AMPARs. These data define unique extracellular interactions with AMPAR complexes and their potential role in coordinating excitatory synaptic plasticity.

## Materials and Methods

### Antibodies and reagents

BioSITe-proteomics: Anti-biotin antibody 1 (Abcam, no. ab53494), anti-biotin antibody 2 (Bethyl Laboratories, no. 150-109A), streptavidin-HRP (Abcam, no. ab7403), protein G beads (EMD Millipore, 760 no. 16-266), high-capacity NeutrAvidin agarose (Thermo Fisher Scientific, no. 29202), biotin (Sigma-Aldrich, no. B4501), biotin-2′,2′,3′,3′-d4 (Sigma-Aldrich, no. 809068), Lipofectamine 2000 (Thermo Fisher Scientific, no. 11668019), biotin-phenol (Iris Biotech, no. CDX-B0270-M100), sequencing-grade trypsin (Promega, no. V5113), GalT1 enzymatic labeling kit (Invitrogen, no. C33368), PNGase F (New England Biolabs, no. P0704S), Click-IT Biotin DIBO Alkyne (Thermo Fisher Scientific, no. C10412), and trypsin (Worthington Biochemical Corporation, no. LS003741) were used. AMPAR antibodies: GluA1 c-terminal (rabbit pAb, clone #4294, homemade), GluA2 N-terminal (clone 19.1, mouse IgG2b, homemade)

### Cloning and molecular biology

The sequence of all constructs used were confirmed with DNA sequencing. All overexpression constructs used were subcloned into the pCAG backbone. CRISPR KO plasmids were subcloned into pU6-(Bbsl)-CBh-Cas9-T2A-mCherry (Addgene #64324). All AMPA receptor subunits use rat cDNAs.

#### APEX2 plasmids

Myc-APEX2 was subcloned into the N-terminus of rat GluA2 flip cDNA in pCAG backbone after the endogenous signal peptide and flanked by short linkers. HA-GluA1/pCAG was generated by subcloning HA into the N-terminus of rat GluA1 flop cDNA after the endogenous signal peptide, flanked by linkers.

AMPA receptor subunits were rat cDNAs, GluA1 flop and GluA2 flip variants. OBCAM and NTM mouse cDNAs were used.

#### CRISPR KO

CRISPR KO guides targeting OBCAM and NTM were subcloned into pU6-(Bbsl)-CBh-Cas9-T2A-mCherry (Addgene #64324). KO guide sequences:

OBCAM KO guide #1: caccgTTGACCAAGATGATCACTCG

OBCAM KO guide #2: caccgACAGGTGTACCATAGATGAC

NTM KO guide #1: caccgTGACAACCGAGTCACCCGGG

NTM KO guide #2: caccGTGGTGCCTAGATCCTCGTG

## ANIMALS

Sprague Dawley rats were used to generate embryos for neuronal cultures. All animals were treated in accordance with the Johns Hopkins University Animal Care and Use Committee guidelines.

### Neuronal Culture

Neuronal cultures were generated as previously described^50,69^. Timed pregnant Sprague-Dawley rats were purchased and dissected at embryonic day 17 or 18. Cortical cells were plated onto 10cm dishes coated with 1 mg/ml Poly-L-Lysine hydrobromide (Sigma) at a density of 4e6/dish and grown in Neurobasal Plus Medium (Gibco) supplemented with 2% B-27 Plus (Gibco), 2mM GlutaMAX (Thermo Fisher Scientific), Penicillin-Streptomycin (100U/ml, Thermo Fisher Scientific), and 5% horse serum (Hyclone). FDU (Sigma) was added at days in vitro (DIV) 4 in NM1 (Neurobasal Plus Medium, 2% B-27 Plus, 2mM GlutaMAX, Penicillin-Streptomycin (100U/ml), 1% horse serum). Neurons were subsequently fed two times per week until experiments were performed with serum-free media (NM0, Neurobasal Plus Medium, 2% B-27 Plus, 2mM GlutaMAX, Penicillin-Streptomycin (100U/ml). Hippocampal cells were plated at a density of 100K/well (of a 12-well plate) in the same manner except with standard Neurobasal Medium and standard B27. For hippocampal cultures, cells were swapped to serum-free media at DIV1 and subsequently fed once a week with serum-free media.

For immunostaining experiments, hippocampal neurons were transfected at DIV 17-18 using lipofectamine 2000 (Invitrogen) according to manufacturer’s instruction. Cells were fixed and stained 48hr after transfection. For FRAP experiments, hippocampal neurons were transfected at DIV 11-12 using lipofectamine 2000 (Invitrogen) according to manufacturer’s instruction. SEP-GluA2 live-cell imaging was performed DIV 19-21. For APEX2 labeling experiments (mass spectrometry or biochemical analysis), dissociated cortical cultures were electroporated with APEX2-GluA2, HA-GluA1, and dsRed (cell fill) at DIV 0 using Rat Neuron Nucleofector kit according to manufacturer’s protocol (Lonza Group)^70^.

### HEK 293 cells

HEK 293T cells (ATCC: CRL-3216^™^) were grown on 10cm plates (for biochemistry) or on collagen (Advanced Biomatrix) coated glass coverslips in 12-well plates (for immunofluorescence and live-cell imaging) in DMEM (Gibco) supplemented with 10% Fetal Bovine Serum (Hyclone) and Penicillin-Streptomycin antibiotics (100 U/ml, Thermo Fisher Scientific). Cells were maintained at 37C with 5% CO2. For biochemistry (GST-Pulldown experiments) HEK cells were transfected via Calcium Phosphate precipitation. For immunofluorescence and live-cell imaging experiments HEK cells were transfected with Lipofectamine 2000 (Invitrogen).

### PSD isolation from cultured neurons

P2 and PSD fractions were isolated from cultured rat cortical neurons as described previously^69,70^. Briefly, mature rat cortical neurons (DIV 19-21) were harvested in homogenization buffer (320 mM sucrose, 10 mM Hepes pH 7.4, 5 mM sodium pyrophosphate, 1 mM EDTA, protease inhibitor mixture (Roche), 200 nM okadaic acid), and homogenized using a 26.5-gauge needle. Homogenate was centrifuged at 1000 × g for 10 min at 4°C yielding P1 and S1. S1 was centrifuged at 17,000 × g for 20 min at 4°C to yield P2 and S2. P2 was resuspended in water and quickly adjusted to 4 mM Hepes pH 7.4, followed by 30-min agitation at 4°C. The lysed P2 fraction collected by centrifugation at 25,000 × g for 20 min at 4°C. The resulting pellet was resuspended in 50 mM Hepes pH 7.4 and an equal volume of 1% triton X-100, and agitated for 10 min 4°C. PSD fractions were obtained by centrifugation at 32,000 × g for 20 min at 4°C. The final PSD pellet was resuspended in 1%NP40, 0.5%DOC, 0.1%SDS lysis buffer and sonicated. Protein quantification was performed via BCA assay, samples were diluted in Laemmli Buffer (BioRad) containing 5% β-mercaptoethanol and frozen until analyzed by western blotting.

### Immunocytochemistry

Neurons were rinsed with PBS and fixed for 15 minutes at room temperature in PBS containing 4% paraformaldehyde (PFA) and 4% sucrose. Cells were washed 4 times with PBS and were incubated with primary antibodies overnight at 4°C in 1x GDB buffer (30 mM phosphate buffer (pH 7.4) containing 0.2% gelatin, 0.3% Triton X-100, and 0.25 M NaCl). Cells were washed 5 times with 1x PBS and incubated with secondary antibodies (1:500) for 1h at room temperature in the dark (goat-conjugated Alexa Fluor conjugated 647, 568, or 488, Thermo Fisher Scientific). Cells were washed 5 times with PBS and then mounted onto glass slides with Fluoromount-G (Southern Biotech). Images were acquired on a confocal microscope (Zeiss LSM 880) at optimal resolution with a 63x objective.

### Glycine-induced chemical LTP (cLTP)

Rat cortical neurons (DIV 17-18) were treated with 200uM AP5 in culture media. 48hrs later, cells were incubated for 2.5h at 37°C in buffer A, Mg-ACSF (143mM NaCl, 5mM KCl, 1mM MgCl_2_, 2mM CaCl2, 10mM Glucose, 10mM Hepes, 100 μM PTX, pH 7.4, 303 mOsm) containing 1 μM TTX, 100 μM PTX, 1 μM Strychnine, 200 μM AP5. For cLTP stimulation, cells were washed 1x with buffer B (0 Mg ACSF containing 200 μM glycine) and stimulated with buffer B for 10min at 37°C. Cells were then washed 1x with buffer C (buffer A with no AP5) and then recovered in buffer C for 15 minutes at 37°C. For control neurons, all steps and buffer exchanges were performed with buffer A. For cLTP/AP5 control neurons, 200 μM AP5 was included in all buffers (A, B, and C) throughout the experiment.

### BxxP synthesis

BxxP was synthesized as previously described^38^ and yielded **853 mg BxxP (87.5% yield).** LC-MS and NMR were performed to check final product formation, structure, purity.

### APEX labeling with BxxP

APEX labeling was performed by modifying the protocol described previously^38^. For proteomic experiments, BxxP labeling was initiated 15 minutes after the 10 minute cLTP stimulation. For labeling reactions, cells were incubated for 1 minute at room temperature with 300uM BxxP, 1mM H_2_O_2_ in buffer C (cLTP neurons) or buffer A (control and cLTP/AP5 neurons). The reaction was rapidly quenched by washing 3x with ice-cold quenching solution (Mg-ACSF containing 10 mM sodium azide (VWR), 10 mM sodium ascorbate (VWR), and 5 mM Trolox (Sigma)) on ice. Neurons were harvested by gentle scraping in ice-cold homogenization buffer (above) containing protease inhibitors and quenching reagents (10 mM sodium azide, 10 mM sodium ascorbate, and 5 mM Trolox. Cells were pelleted by centrifugation at 3000 xg for 10 minutes at 4°C. The supernatant was discarded and pellets were stored at −80°C. For western blotting and streptavidin-pull down experiments, control neurons were used. For proteomics experiments, N=1 for each condition contained 10-10cm dishes of labeled cortical neurons from 1-3 independent labeling experiments and batches of neurons.

Replicates were subsequently used in the 3 independent BioSITe-proteomics experiments which were ultimately performed.

### BioSITe Mass Spectrometry

Samples were prepared for BioSITe-based proteomic analysis as described previously^48,51^.

### Peptide preparation

For each replicate, 10 mg of total protein from each condition (control, cLTP and cLTP/APV) were digested into peptides. Total protein amount was estimated by cell number (4e6 per 10cm dish, 1-10cm dish = 1mg protein). Briefly, cell pellets were resuspended in 8M Urea/50mM TEABC buffer with protease inhibitors, sonicated (three rounds, duty cycle 30%, 20 s pulses) and centrifuged 16,000 xg 10 min at 4°C. Supernatants were then transferred to fresh tubes, DTT was added to a final concentration of 10 mM and incubated at 37°C for 1 hr to fully denature proteins prior to digestion. Samples were cooled to room temperature, and IAA was added to a final concentration of 30 mM and incubated for 1 hr in the dark. Lysates were then diluted 4x with 50mM TEABC buffer to dilute the urea concentraation to <2M, and 100 μg trypsin was added to each mg of total protein at 1:10 enzyme to protein ratio. Lysates were digested overnight at 37°C and checked by SDS-PAGE to confirm complete digestion. Peptides were then acidified by adding TFA to a final concentration of 1% and incubated at room temperature for 10 min. Peptides were then centrifuged at 10,000 xg for 10 min at room temperature and supernatant containing peptides was transferred to a fresh tube. The resulting tryptic peptides were desalted using a Sep-PAK C_18_ (Waters Corp.) and subsequently lyophilized.

### BioSITe

The Protein G beads (EMD Millipore) were 3x washed with PBS. Anti-biotin antibodies (Bethyl Laboratories) were coupled to 120 μL of protein G bead slurry overnight at 4°C. Antibody-coupled beads were further washed with PBS once and BioSITe capture buffer (50 mM Tris, 150 mM NaCl, 0.5% Triton X-100) twice. Peptides were then dissolved in 1 mL of BioSITe capture buffer. After dissolving peptides, pH was adjusted to 7.0 to 7.5 and peptide BCA was performed to estimate peptide concentration. 8 mg peptides were used per replicate. Peptides were subsequently incubated with anti-biotin antibody-bound protein G beads for 1 hr at 4°C. The bead slurry was sequentially washed three times with BioSITe capture buffer, three times with 50 mM Tris-HCl, pH 7.5, and three times with ultrapure water. Biotinylated peptides were then eluted four times using elution buffer (80% acetonitrile and 0.2% trifluoroacetic acid in water). The eluent was further cleaned up using C_18_ reversed-phase column and then analyzed on an Orbitrap Fusion Lumos mass spectrometer.

### Mass Spectrometric Analysis and Post-Processing and Bioinformatics were performed as described previously ^6,7^

Briefly, Proteome Discoverer (v2.1; Thermo Scientific) suite was used for quantitation and identification using all three replicate LC−MS/MS runs per experiment searched together. Spectrum selector was used to import spectrum from raw file. During MS/MS preprocessing, the top 10 peaks in each window of 100 m/z were selected for database search. The tandem mass spectrometry data were then searched using SEQUEST algorithm against protein databases with the addition of Fasta file entries for APEX2-GluA2 and HA-GluA1. The search parameters for identification of biotinylated peptides were as follows: (a) trypsin as a proteolytic enzyme (with up to three missed cleavages); (b) peptide mass error tolerance of 10 ppm; (c) fragment mass error tolerance of 0.02 Da; and (d) carbamidomethylation of cysteine (+57.02146 Da) as a fixed modification and oxidation of methionine (+15.99492 Da) and covalent binding of BxxP(+587.31414 Da), Oxidized BxxP (+603.3091 Da) and their +1 isotopologues (+588.3175 Da and 604.31241 Da) to tyrosine as variable modifications.

The data were analyzed in Proteome Discoverer (Thermo Scientific, v2.2.0.388) using a tiered search strategy. First, the m/z values of precursor and fragment ions were recalibrated using the “Spectrum Filers RC” module to adjust for drifts in instrument mass accuracy. The recalibrated data were then analyzed by MASCOT (Matrix Science, v2.6.2) against the SwissProt *Rattus* protein database (Mention total number of protein sequence entries e.g. 20,000 fasta sequences) using mass tolerances of 6 ppm and 0.01 Da for precursor and fragment ions, respectively. Deamidation of asparagine and glutamine and oxidation of methionine residues were set as dynamic modifications and carbamidomethylation of cysteine was set as a static modification. Percolator 3.00 was used to score the peptide spectral matches. The first tier was intended to filter out spectra which could be easily attributed to *Rattus* proteins. Spectra which did not meet the 1% FDR threshold of the first search were analyzed with the Byonic algorithm against the same Rattus protein database appended with the sequences of recombinant APEX2 and GluA proteins. The search used 6 ppm tolerances for both precursor and fragment ions and included the following modifications with up to 2 dynamic modifications allowed per peptide:

-Static carbamidomethylation of all cysteines

-Common1 tier: dynamic offsite carbamidomethylation of aspartic acid, glutamic acid, histidine, lysine, serine, threonine, and tyrosine residues, deamidation of asparagine and glutamine, and oxidation of methionine.

-Common2 tier: acetylation of the N terminus and lysine residues, and pyro-glutamation of N terminal glutamine.

-Rare tier: lysine and arginine carbamylation; phosphorylation of serine threonine and tyrosine; trioxidation of tryptophan; formylation of N-terminus, lysine, serine and threonine; dimethylation of lysine and arginine, and covalent binding of BxxP(+587.31414 Da), Oxidized BxxP (+603.3091 Da) and their +1 isotopologues (+588.3175 Da and 604.31241 Da) to tyrosine. Percolator was again used to score peptide spectra matches and a 1% FDR cut-off was applied. Modifications identified by Byonic were localized using the ptmRS module and all identified peptides were quantified by integrated precursor ion peak area using the Minora Feature Detector node.

### Coimmunoprecipitation (coIP)

*HEK cells.* Cells were seeded in 6 well dishes for coIP experiments, and 1 well of confluent, transfected cells was used for 1 coIP. HEK cells were lysed in coIP buffer (PBS containing 50 mM NaF, 5 mM sodium pyrophosphate, 1% Nonidet P-40, 0.5% sodium deoxycholate, 1 μM okadaic acid, 2.5 mM sodium orthovanadate, and protease inhibitor mixture (Roche)), solubilized for 1hr at 4°C and then centrifuged at 17,000 xg for 10 minutes at 4°C. The resulting lysates were incubated with anti-Myc antibodies (mouse IgG1, homemade) overnight at 4°C. 15μL Protein G beads were added to each coIP the following day for 1h at 4°C, and then beads were washed 5x with coIP buffer and then eluted in 2x sample buffer (BioRad) containing 5% β-mercaptoethanol. Bound proteins were resolved by SDS-polyacrylamide gel electrophoresis for Western blot analysis.

#### Cortical neurons

The P2 membrane fraction was lysed in coIP buffer (PBS containing 50 mM NaF, 5 mM sodium pyrophosphate, 1% Nonidet P-40, 0.5% sodium deoxycholate, 1 μM okadaic acid, 2.5 mM sodium orthovanadate, and protease inhibitor mixture (Roche)). Pellets were solubilized for 1hr at 4°C and then centrifuged at 17,000 xg for 10 minutes at 4°C. The P2 lysates were transferred to new tubes and protein concentration was measured via BCA and 200μg P2 lysate was used for each coIP. P2 lysates were incubated with AMPAR antibodies overnight at 4°C in 300μL total volume. 15μL Protein G beads were added to each coIP the following day for 1h at 4°C, and then beads were washed 5x with coIP buffer and then eluted in 2x sample buffer (BioRad) containing 5% β-mercaptoethanol. Antibodies used for coIPs: GluA1 C-terminal (Rabbit 4294, homemade), GluA2 N-terminal (Cell Signaling, rabbit mAb, clone E1L8U), GluA2 C-terminal (NeuroMAb, clone L21/32). Bound proteins were resolved by SDS-polyacrylamide gel electrophoresis for western blot analysis.

### Streptavidin pull-down

Labeled cell pellets were thawed in ice-cold coIP buffer (PBS containing 50 mM NaF, 5 mM sodium pyrophosphate, 1% Nonidet P-40, 0.5% sodium deoxycholate, 1 μM okadaic acid, 2.5 mM sodium orthovanadate, and protease inhibitor mixture (Roche)). Cells were solubilized for 2h at 4°C, and lysates were cleared by centrifugation at 17,000 xg for 20min. For streptavidin pull-downs, 250μg labeled neuronal lysates were mixed with 75μL streptavidin magnetic beads (Thermo Fisher #88817) that had been washed 3x. The total volume was brought to 600 μL and 10mM DTT (final concentration) was added to each pull-down before they were incubated overnight at 4°C. Beads were then vigorously washed with 1mL of each of the following (in order): RIPA, 1M KCl, 2%SDS in 50mM Tris-HCl, pH7.4 (room temperature wash), RIPA (2 final washes). Biotinylated proteins were then eluted from the beads using pre-warmed 2x SDS sample buffer (4mM biotin, 40mM DTT, 5% β-ME) by heating for 20 minutes at 100°C. Proteins were quickly removed from the beads immediately after removal from the heat block and frozen until they were analyzed western blot analysis.

### FRAP Imaging and analysis

All confocal images were acquired with a Zeiss LSM 880 microscope. Neurons were imaged at 37°C in Mg-ACSF (143mM NaCl, 5mM KCl, 1mM MgCl_2_, 2mM CaCl2, 10mM Glucose, 10mM Hepes, 100 μM PTX, pH 7.4, 303 mOsm) in a humidity-controlled chamber. The laser power and repetitions needed for successful bleaching was optimized and kept consistent across experiments. Images were collected with oil-immersive 63x objective. Time-series Z-stacks were acquired with acquired every 20 seconds for 40 cycles, with 2 time points before photobleaching. For each dendritic region at least 5 putative synaptic puncta (selected by neuronal morphology, on spines on secondary dendrites) were bleached, ensuring that there were several puncta in the field of view that were not bleached. Images were analyzed in FIJI. Maximum image projections were generated for the time series, and a median filter was applied to all channels (1-pixel) and the mCherry cell-fill channel was used to correct for movement during imaging with a rigid-body transformation (MultiStackReg plugin). Regions of interest (ROIs) were drawn around bleached and unbleached puncta (5 per image) as well as regions of background (away from the dendritic signal,3 per image). Background signal was subtracted from the puncta signal, which was then normalized to the average intensity of the unbleached puncta at each time point (to account for the low levels of acquisition bleach over time). The signal in the bleached ROIs was then normalized such that the average baseline was centered on 1 and the post-bleach time point was 0. For each puncta the estimated maximum recovery was estimated with a one phase exponential fit (in GraphPad Prism 9).

## Acknowledgments

We would like to thank all the members of the R.L.H. lab for helpful discussion and critical reading of the manuscript, especially Richard Johnson for experimental support. This work was supported by the NIH (F32 NS098946) H.G.M., (K99 MH132811) W.D.H., (ROO MH124920) A.M.B. and (R37 NS036715) R.L.H.

