## Supplemental Figures for "Dynamic extracellular interactions with AMPA receptors"

**SUPPLEMENTAL FIGURE LEGENDS**

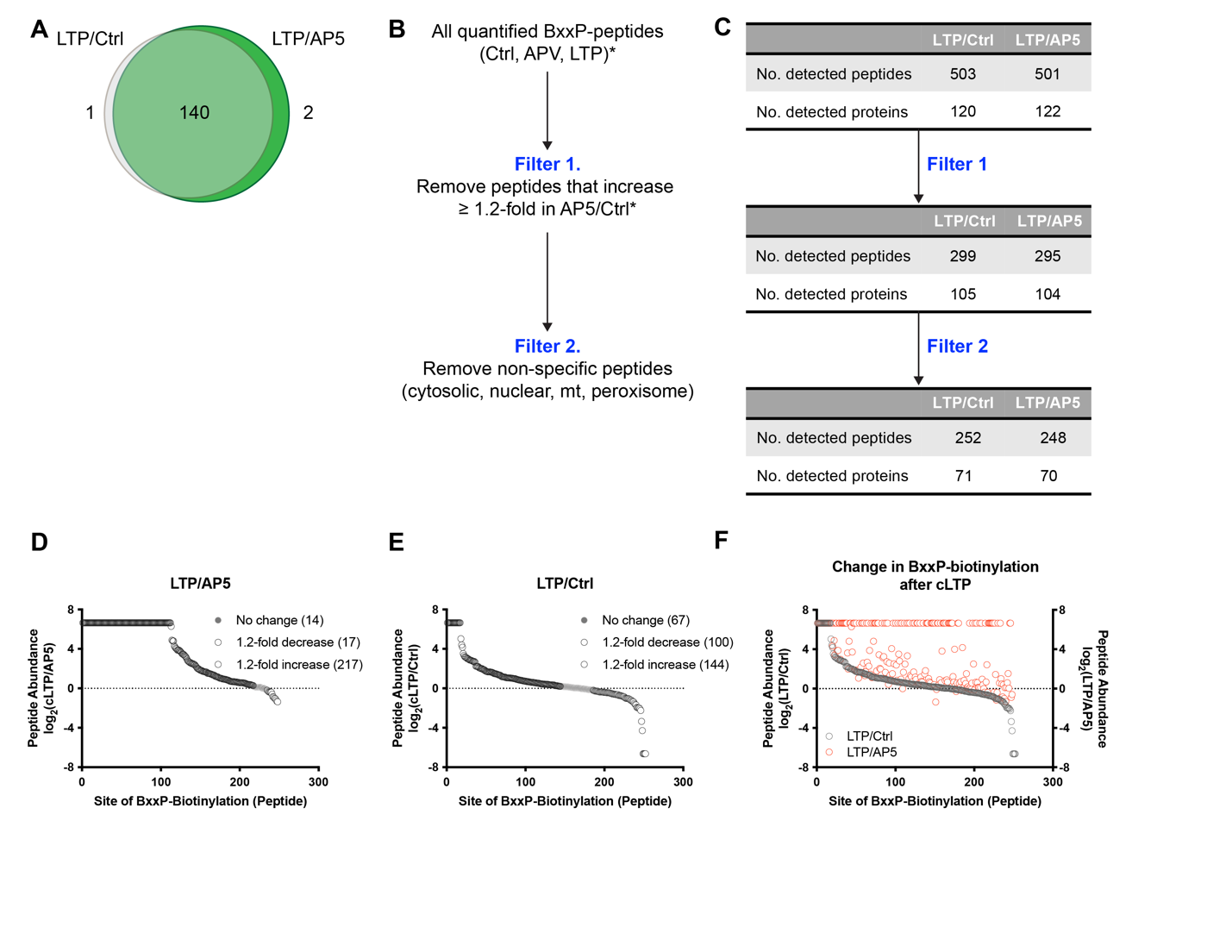

**SUPPLEMENTAL FIGURE 1. Differential biotinylation by APEX2-GluA2 AMPARs after cLTP**

**(A)** Venn diagram showing the overlap of proteins identified by BioSITe-MS. Ratios of differential biotinylation of individual peptides between LTP and Ctrl or AP5 conditions were calculated. To determine the overlap across these 2 groups, we compared the proteins identified between these two groups (Venn diagram) that represent all peptides that were quantified in either LTP/Ctrl or LTP/AP5. **(B-C)** All peptides identified by BioSITe-MS were filtered to remove peptides that did not change in abundance after cLTP (filter 1) and non-specific proteins (filter 2). The number of peptides and proteins after each filter was applied are shown in supplemental Figure 2C. **(D-F)** Differential biotinylation after cLTP by APEX2-AMPARs. Graphs showing the change in abundance of all peptides remaining after filters 1 and 2 were applied to the raw data. **(D)** LTP/AP5 and **(E)** LTP/Ctrl, number of peptides that do not change, increase or decrease after cLTP are shown. **(F)** Peptide abundances calculated from LTP/Ctrl and LTP/AP5 (red) samples are shown for individual peptides (x-axis).

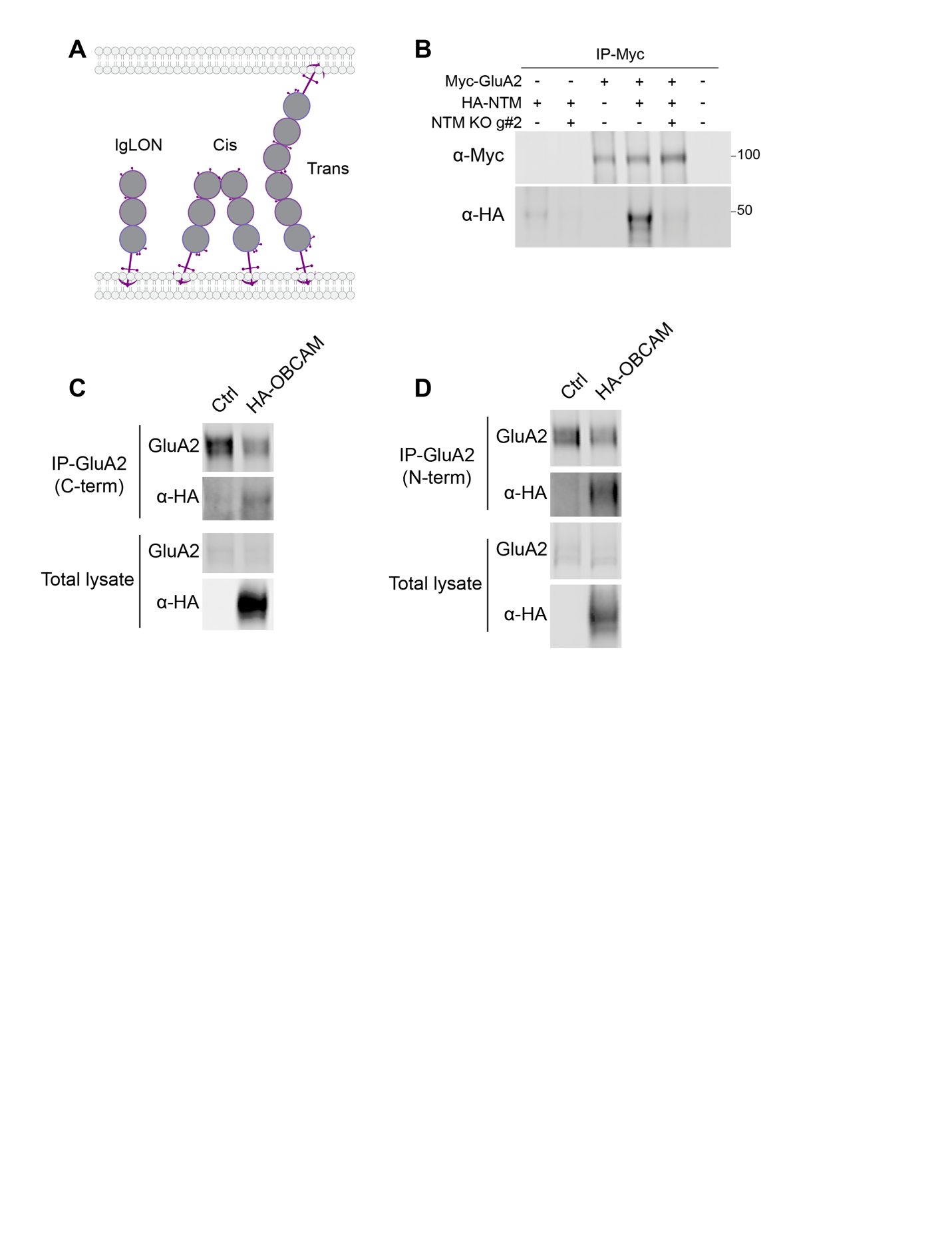

**SUPPLEMENTAL FIGURE 2. IgLONs interact with the extracellular region of AMPARs**

**(A)** Schematic of IgLON proteins. IgLONs consist of 3-Ig domains (grey circle) anchored to the extracellular side of the plasma membrane via GPI-anchor (purple). IgLONs are know to interact with themselves or other IgLON proteins through the third Ig domain (ref) in cis (same side of the membrane) or trans (interaction occurs between IgLONs residing on two different membranes). **(B)** NTM directly interacts with GluA2-containing AMPARs. coIP experiments in HEK cells show HA-NTM is present in GluA2 precipitates (lane 4). To demonstrate specificity of this interaction we used a CRISPR KO guide targeting NTM cDNA. Expression of NTM is successfully depleted by expression of CRISPR knockout guides targeting the cDNA (NTM KO g#2) (lane 2) and therefore depletes NTM from GluA2 precipitates (lane 5). **(C-D)** IgLONs interact with endogenous AMPAR complexes in neurons. Neurons expressing HA-tagged OBCAM were fractionated and endogenous AMPARs were purified from crude membrane fractions using anti-GluA2 antibodies targeting the intracellular C-terminus (C) or extracellular N-epitopes on GluA2 (D). Immunoblotting with anti-HA antibodies shows HA-OBCAM precipitate with GluA2-containing AMPARs, but not in control neurons.

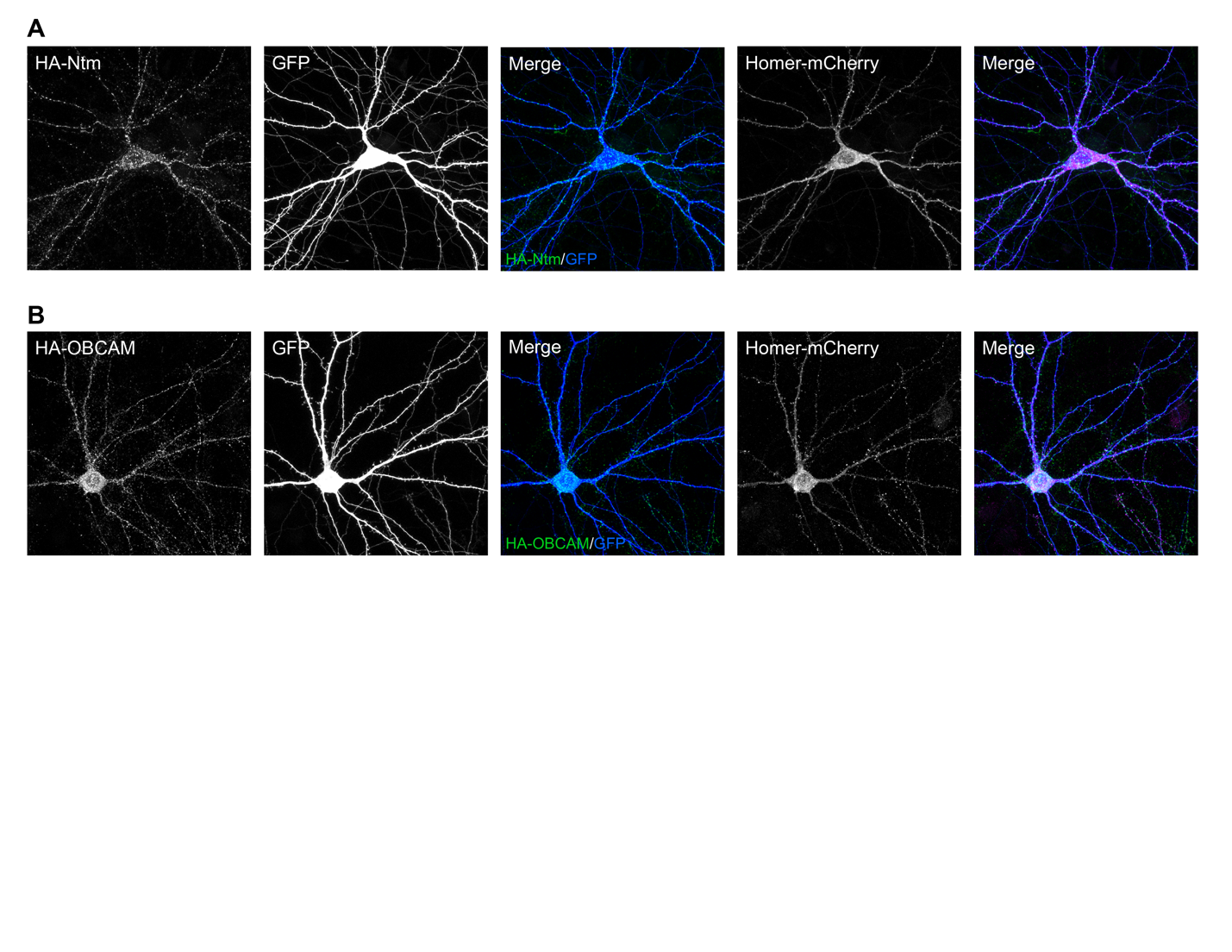

**SUPPLEMENTAL FIGURE 3. Overexpressed NTM and OBCAM show synaptic localization. (A)** Cultured rat neuron transfected with HA-tagged NTM (far left) showing a distinct punctate distribution compared to the GFP cell fill (middle left). NTM puncta show partial overlap with Homer-mCherry puncta (middle right, far right merge). **(B)** Cultured rat neuron transfected with HA-tagged OBCAM (far left) showing a similar distribution and similar overlap with Homer-mCherry (middle right, far right merge).

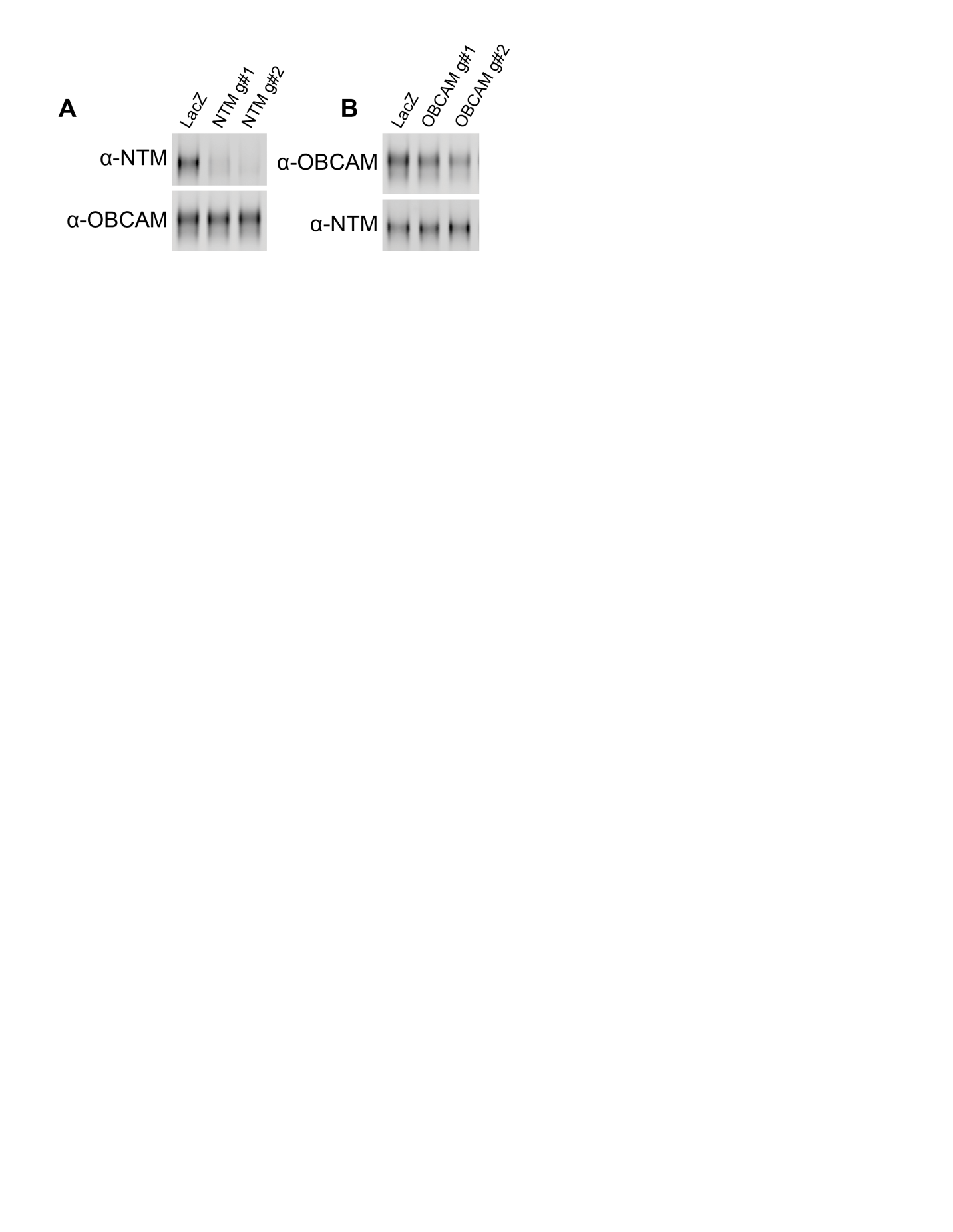

**SUPPLEMENTAL FIGURE 4. CRISPR KO of NTM and OBCAM in neurons.** Cultured rat neurons electroporated with different CRISPR KO guides (#1, #2) against NTM **(A)** or OBCAM **(B)**. Western blotting for NTM and OBCAM shows each KO guide is selective for the targeted IgLON, and does not alter protein levels of other IgLONs.
